# wg-blimp: an end-to-end analysis pipeline for whole genome bisulfite sequencing data

**DOI:** 10.1101/859900

**Authors:** Marius Wöste, Elsa Leitão, Sandra Laurentino, Bernhard Horsthemke, Sven Rahmann, Christopher Schröder

## Abstract

**Background:** Analysing whole genome bisulfite sequencing datasets is a data-intensive task that requires comprehensive and reproducible workflows to generate valid results. While many algorithms have been developed for tasks such as alignment, comprehensive end-to-end pipelines are still sparse. Furthermore, previous pipelines lack features or show technical deficiencies, thus impeding analyses.

**Results:** We developed wg-blimp (<u>w</u>hole <u>g</u>enome <u>b</u>isulfite sequencing <u>m</u>ethylation analysis <u>p</u>ipeline) as an end-to-end pipeline to ease whole genome bisulfite sequencing data analysis. It integrates established algorithms for alignment, quality control, methylation calling, detection of differentially methylated regions, and methylome segmentation, requiring only a reference genome and raw sequencing data as input. Comparing wg-blimp to previous end-to-end pipelines reveals similar setups for common sequence processing tasks, but shows differences for post-alignment analyses. We improve on previous pipelines by providing a more comprehensive analysis workflow as well as an interactive user interface. To demonstrate wg-blimp’s ability to produce correct results we used it to call differentially methylated regions for two publicly available datasets. We were able to replicate 112 of 114 previously published regions, and found results to be consistent with previous findings. We further applied wg-blimp to a publicly available sample of embryonic stem cells to showcase methylome segmentation. As expected, unmethylated regions were in close proximity of transcription start sites. Segmentation results were consistent with previous analyses, despite different reference genomes and sequencing techniques.

**Conclusions:** wg-blimp provides a comprehensive analysis pipeline for whole genome bisulfite sequencing data as well as a user interface for simplified result inspection. We demonstrated its applicability by analysing multiple publicly available datasets. Thus, wg-blimp is a relevant alternative to previous analysis pipelines and may facilitate future epigenetic research.

## Background

Since the development of DNA sequencing, a large number of studies on genetic variation have been conducted, while extensive research on the epigenetic level has only emerged in the recent past. Although most cells within an organism are identical in their genomic sequence, different tissues and cell types vary in their patterns of epigenetic modifications that confer their particular identity. DNA methylation is one of the most important epigenetic marks and occurs mainly at CpG dinucleotides. There are almost 28 million of such sites in the human genome, thus 450k arrays (which cover only 1.6% of all CpGs) are not sufficient to detect small differentially methylated regions (DMRs) (1). As a result, data-intensive whole genome bisulfite sequencing (WGBS) is required to properly identify all CpG methylation levels. While the costs for generating these data sets have been very high, the continuous and sustained reduction of sequencing costs allows more and more WGBS datasets to be generated, creating the need for comprehensive and reproducible analysis tools. Many algorithms have already been established for different aspects of WGBS analyses such as alignment and DMR detection. However, choosing appropriate algorithms and integrating them into an end-to-end analysis workflow is not a trivial task due to combinatorial explosion of possible pipeline setups. Setting up an end-to-end WGBS analysis workflow is further hindered by different requirements of interacting tools, e.g. input and output formats or chromosome naming conventions. Previously developed end-to-end pipelines already consider these problems and only require users to supply their raw data and configuration. However, we find previous approaches to lack features required in common research settings, e.g. methylome segmentation, as well as technical limitations such as installation issues, as described in more detail in the *Results & Discussion* section. As a result, we developed a pipeline approach to address these issues.

## Implementation

We present here wg-blimp (<u>w</u>hole <u>g</u>enome <u>b</u>isulfite sequencing <u>m</u>ethylation analysis <u>p</u>ipeline), a workflow for automated in silico processing of WGBS data. It consists of a comprehensive WGBS data analysis pipeline as well as a user interface for simplified inspection of datasets and potential sharing of results with other researchers. Figure 1 gives an overview of the analysis steps provided.

**Fig. 1.**
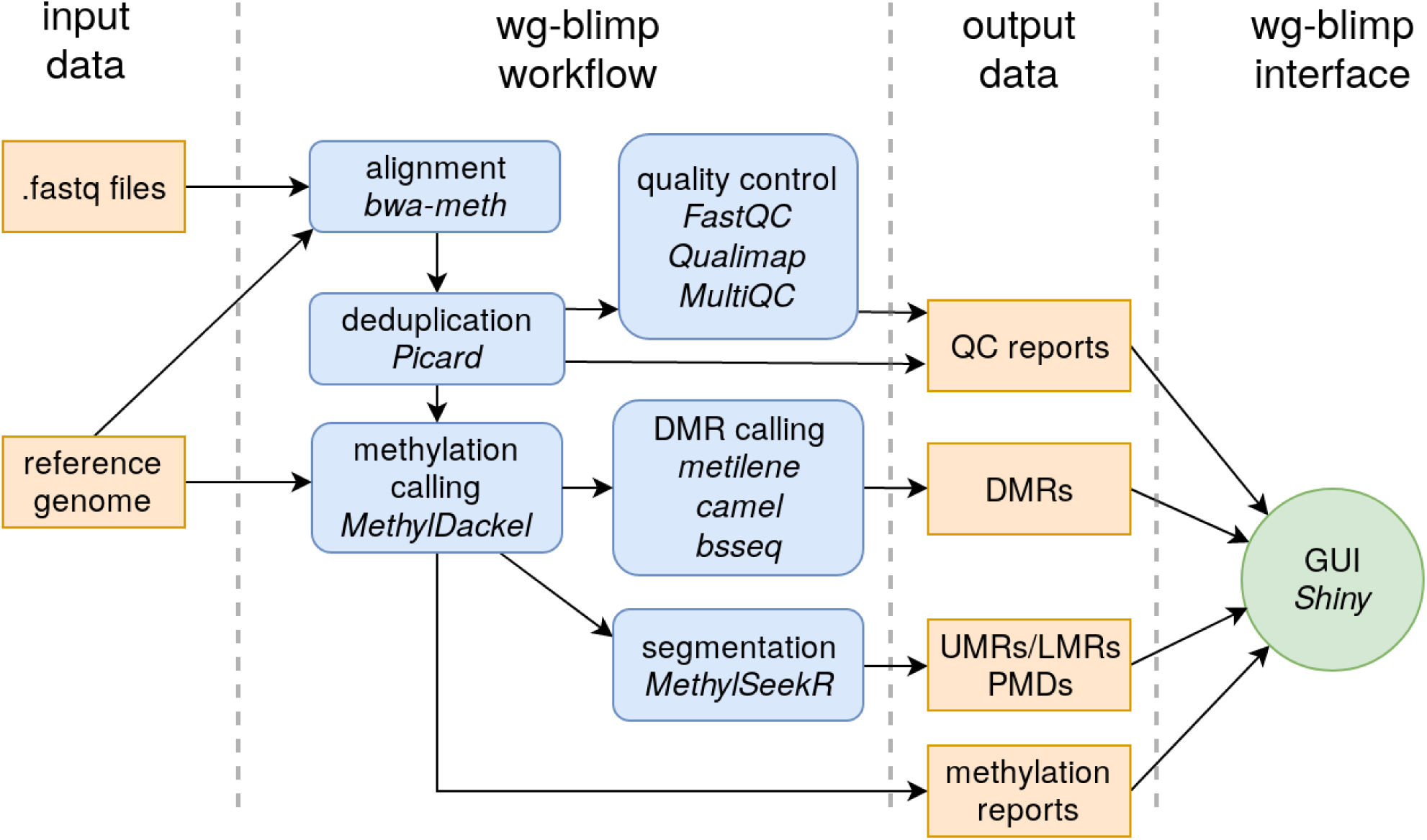
wg-blimp workflow overview. Users only need to provide FASTQ files and a reference genome, and wg-blimp will perform alignment, deduplication, QC checks, DMR calling, segmentation and annotation. Once the pipeline results are available, users can inspect results using a web interface.

With FASTQ files and a reference genome as input, wg-blimp performs a complete workflow from alignment to DMR analysis, segmentation and annotation. We choose bwa-meth (2) for alignment as it provides efficient and robust mappings due to its internal usage of BWA-MEM (3). We omit pre-alignment trimming of reads because of bwa-meth’s internal usage of soft-clipping to mask non-matching read subsequences. Alignments are deduplicated using the Picard toolkit (4). Methylation calling is performed by MethylDackel (5) as it is the recommended tool for use with bwa-meth. Based on the methylation reports created by MethylDackel, wg-blimp computes global methylation statistics. Computing per-chromosome methylation is optional and enables estimation of C > T conversion rates, as unmethylated lambda DNA is commonly added to genomic DNA prior to bisulfite treatment.

For quality control (QC) we use FastQC (6) to evaluate read quality scores. Coverage reports containing information about overall and per-chromosome coverage are generated by Qualimap (7). Qualimap also reports metrics such as GC content, duplication rate, and clipping profiles, thus enabling in-depth quality evaluation of each sample analysed. Quality reports by Picard, Qualimap and FastQC are aggregated into a single interactive HTML report using MultiQC (8)

Multiple algorithms are supported for DMR calling: metilene (9), bsseq (10) and camel (11) are frequently used tools. The application of more than one DMR calling tool is recommended, because these tools identify different, although overlapping sets of DMRs.

We further integrate detection of unmethylated regions (UMRs) and low-methylated regions (LMRs) to identifify active regulatory regions in an unbiased fashion. This segmentation is implemented using MethylSeekR (12) as it provides automatic inference of model parameters using only a user-defined false-discovery rate (FDR) and methylation cutoff. MethylSeekR also implements detection of regions of highly disordered methylation, termed partially methylated domains (PMDs). The presence of PMDs is influencing UMR/LMR detection and is often unknown a priori. As a result, wg-blimp preemptively performs the MethylSeekR workflow with and without PMD computation. Based on the metrics measured by MethylSeekR users may decide wether or not to consider PMDs when analysing UMRs and LMRs. Resulting DMRs, UMRs, LMRs and PMDs are annotated for overlap with genes, promoters, CpG islands (CGIs) and repetitive elements as reported by Ensembl (13) and UCSC (14) databases. Average coverage per DMR is computed using mosdepth (15) to enable filtering of DMR calls in regions of low coverage.

We base the wg-blimp pipeline on the workflow execution system Snakemake (16) as it enables robust and scalable execution of analysis pipelines and prevents generation of faulty results in case of failure. Snakemake also provides run-time and memory usage logging, thus easing the search for bottlenecks and performance optimization. To minimize errors caused by changing software versions we utilize Bioconda (17) for dependency management and installation.

Once the analysis workflow completes, users may load the results into wg-blimp’s user interface. We implemented the interface using the R Shiny framework that enables seamless integration of R features into a reactive web app. The interface aggregates QC reports, pipeline parameters, and allows inspection and filtering of DMRs based on caller output and annotations. UMRs and LMRs computed by MethylSeekR may also be accessed through wg-blimp’s Shiny interface, and users may dynamically choose whether or not to include PMDs. Since visualization of genomic data is often employed when inspecting analysis results, access links to alignment data for use with the Integrative Genomics Viewer (IGV) (18) are also provided, as IGV provides a bisulfite mode for use with WGBS data.

## Results & Discussion

To evaluate wg-blimp’s relevance for WGBS experiments, we compared it to previous end-to-end pipelines and demonstrated its applicability by analysing three exemplary datasets.

### Comparison to previous pipelines

Since wg-blimp only integrates published software, and exhaustive evaluation of all conceivable pipeline setups would result in combinatorial explosion, we focus here on a feature-wise comparison of pipelines, similar to (19). We compared wg-blimp to BAT (20), bicycle (21), CpG_Me/DMRichR (10, 22–24), ENCODE-DCC’s WGBS pipeline (25), Methy-Pipe (26), Nextflow methylseq (two available workflows) (27), PiGx (28) and snakePipes (19). Pipelines were compared with regards to technical setup (installation, workflow management), WGBS read processing (adapter trimming, alignment, methylation calling, quality control), and post-alignment analyses (DMR detection, segmentation, annotation).

Table 1 gives an overview over each pipeline’s setup. Similar to snakePipes, wg-blimp utilizes Bioconda for installation. Using package managers such as Bioconda or workflow environments like Nextflow (29) not only simplifies installation for users but also provides straightforward update processes of both the pipeline itself as well as its dependencies. Thus, we recommend usage of such package managers to ensure stable runtime environments. For workflow management, we prefer using dedicated workflow management systems such as Snakemake or Nextflow over plain shell scripts, as these allow more scalable and robust execution. Users may also consider using cloud computing platforms such as DNAnexus (dnanexus.com). These platforms alleviate setting up own hardware for analysis, with the downside of users providing their data to third-party providers, thus posing potential data privacy risks.

**Table 1.**
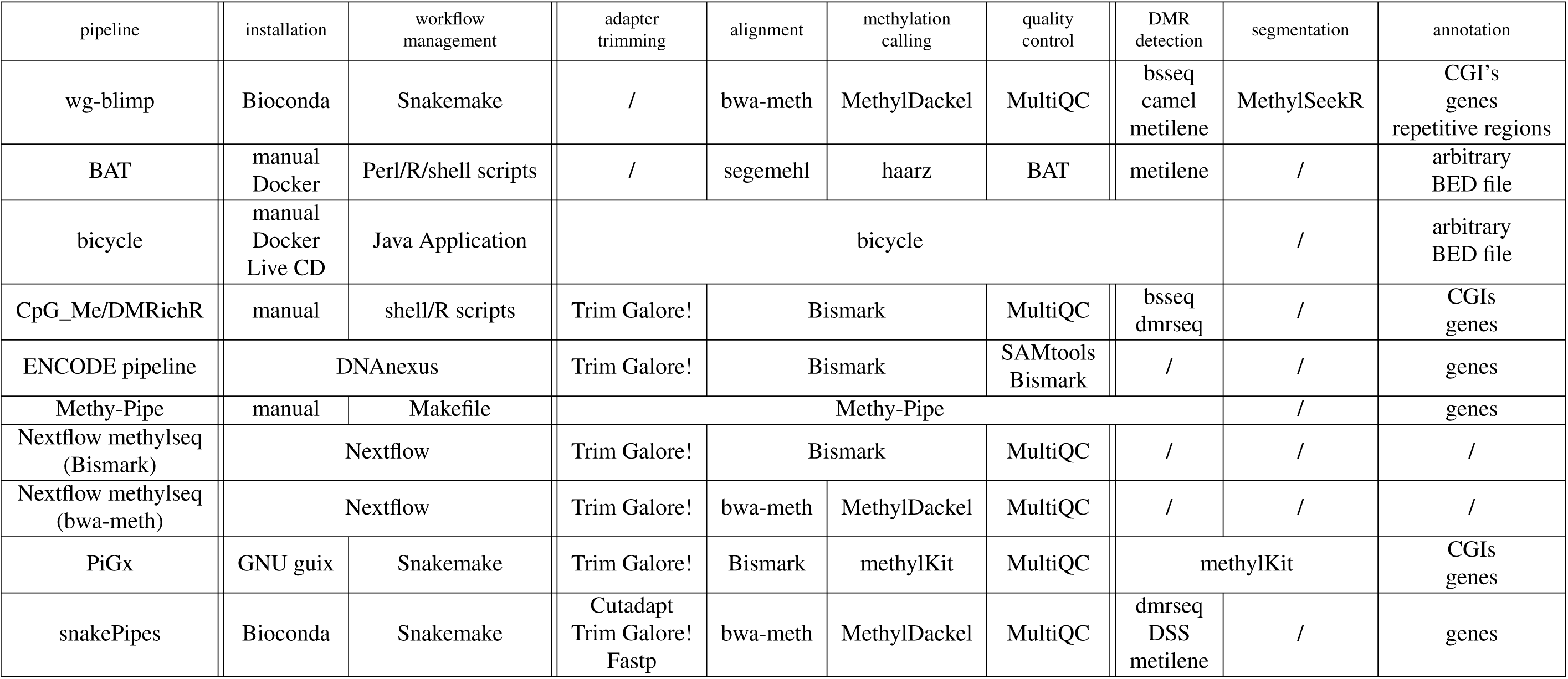
Comparison of WGBS end-to-end pipelines. Most pipelines use similar software for “standard” WGBS analysis tasks such as alignment or QC. wg-blimp improves on existing pipelines by providing a more comprehensive workflow as well as an interactive user interface.

For read processing, wg-blimp employs similar strategies as other pipelines, with popular alignment and methylation calling tools being bwa-meth/MethylDackel and Bismark (24). However, wg-blimp deviates from other pipelines by skipping read trimming, which is handled by BWA-MEM’s soft-clipping. For QC we recommend using MultiQC as it produces HTML quality reports in a compact and scalable way. We omitted the details about which metrics are collected by MultiQC for each pipeline, as the pipelines investigated use common tools such as Picard or sambamba (30) (with the exception of BAT, bicycle and Methy-Pipe).

While most of the pipelines investigated use similar tools for read processing, setups differ for post-alignment analyses. For DMR detection, we pursue a similar setup as snakePipes and BAT by providing multiple DMR callers. wg-blimp and PiGx are the only workflows to perform methylome segmentation. We prefer MethylSeekR over methylKit for segmentation because of its consideration of PMDs.

We further added functionality over other pipelines by implementing an interactive R Shiny GUI. Users may load one or more analysis runs into the Shiny App, thus providing a straightforward way to create a central repository for analysis results to share with fellow researchers. This not only makes distributing individual files unnecessary but also enables a more concise inspection of results. For example, users may switch between segmentation with and without consideration of PMDs using MethylSeekR by toggling a single checkbox instead of having to inspect multiple files. An example of wg-blimp’s interface displaying MethylSeekR results is given in Figure 2. More GUI features are discussed in detail in the Supplementary Material.

**Fig. 2.**
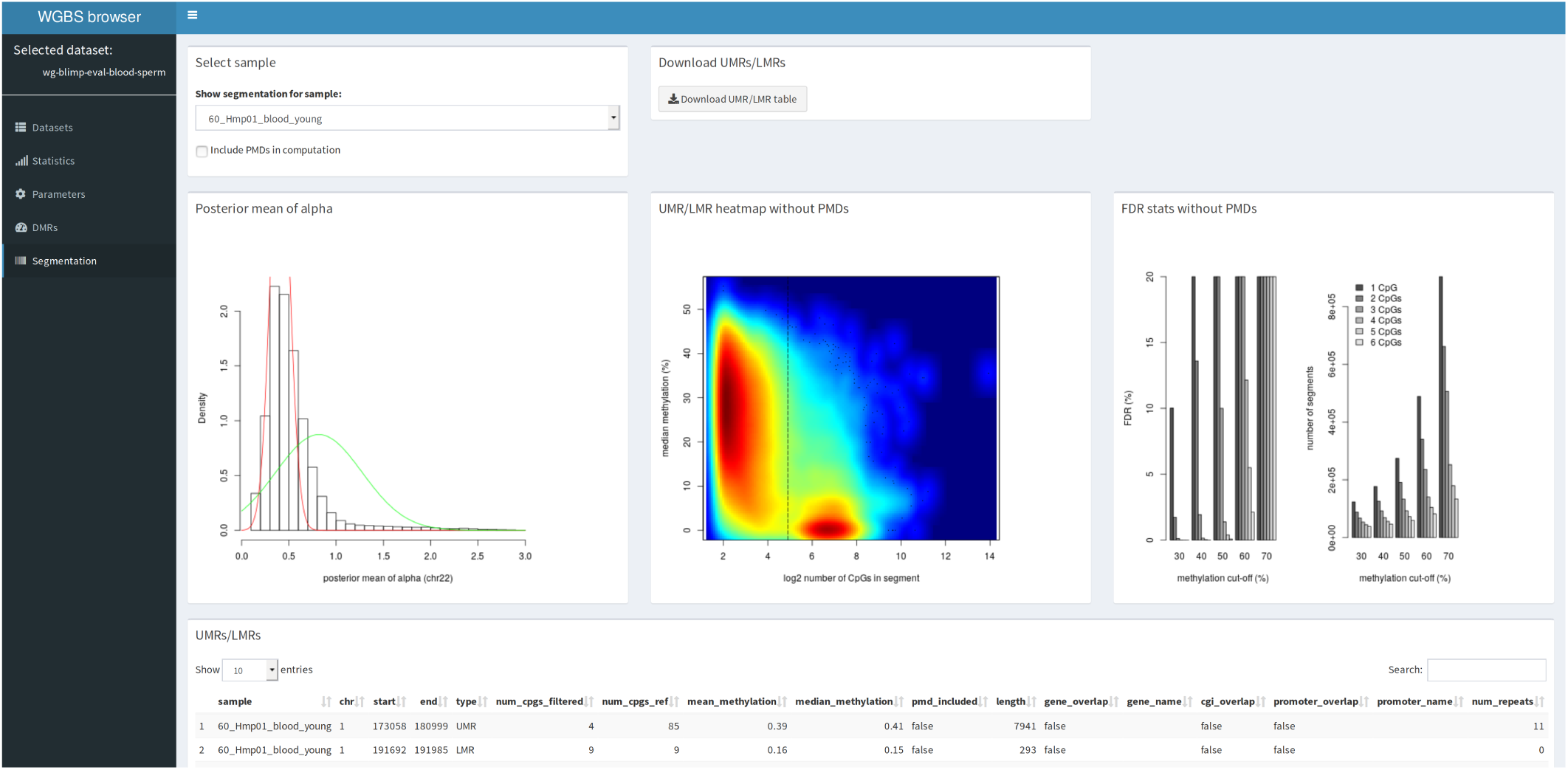
Segmentation tab of wg-blimp R Shiny GUI. Once the analysis pipeline completes, users may load results into wg-blimp’s R Shiny App. The tab depicted here displays MethylSeekR results and allows users to include or exclude PMD computation by toggling a single checkbox.

While we provide additional functionality over previous WGBS pipelines, we would like to emphasize that wg-blimp should not be seen as a replacement for previous approaches, but rather as an extension to the landscape of available workflows. snakePipes, for example, not only provides a WGBS analysis workflow, but is also capable of performing integrative analyses on ChIP-seq, RNA-seq, ATAC-seq, Hi-C and single-cell RNA-seq data. As a result, snakePipes should be preferred over wg-blimp in experiments that aim at integrating different epigenomic assays. In contrast, we prefer wg-blimp over snakePipes for WGBS-only experiments that aim at determining active regulatory regions due to its implementation of segmentation and simplified dataset inspection through its GUI. Thus, when deciding which analysis workflow to choose for a WGBS experiment, we believe there is no “one-fits-all” solution, and we deem wg-blimp one suitable option to consider for future WGBS analyses.

### Application to published datasets

We applied wg-blimp to three exemplary publicly available WGBS datasets. Two of these datasets were utilized to demonstrate wg-blimp’s DMR calling capabilities and a third to demonstrate methylome segmentation. All analyses were executed on a server equipped with two Intel Xeon E5-2695 v4 CPU’s, 528 GB of memory and Debian 9 as operating system (OS). 64 threads were allocated for each analysis.

#### DMR detection

One of the DMR datasets consists of two pairs of isogenic human monocyte and macrophage samples (31), the other of two pairs of isogenic human blood and sperm samples (each generated from pools of DNA from six men) (32). We chose these two datasets to demonstrate wg-blimp’s capability of calling DMRs for cases where few (monocytes vs. macrophages) or many (blood vs. sperm) DMRs are expected due to the degree of relatedness between compared groups.

For the monocyte/macrophage dataset we chose hg38 as reference and used a coverage of at least 5×, at least 4 CpG sites overlapping, and a minimum absolute difference of 0.3 as thresholds for DMR calling. We detected 6,189 DMRs in total, with 4,078 DMRs overlapping genes and 886 DMRs overlapping promoter regions. We were able to recover 112 of the original 114 DMRs reported, even though (31) used hg19 as reference genome and only BSmooth for DMR calling. Most of these DMRs are outside of CpG islands (6,009 DMRs) and lose DNA methylation during differentiation (5,765 DMRs), which is consistent with the original findings (31). Excluding indexing of the reference genome, the whole analysis workflow from FASTQ files to annotated DMRs took 38.87 hours in total. A maximum memory usage of 216.07 GB was reached for bsseq DMR calling (Supplementary Material). bwa-meth alignment was the most time consuming step with a run time of 27.81 hours for a single sample using 16 threads.

For the blood/sperm dataset we used wg-blimp to determine soma-germ cell specific methylation differences. We found 410,247 DMRs (≥ 4 CpGs, ≥ 0.3 absolute difference, ≥ 5× coverage), of which 192,953 overlap with genes, 58,183 with promoters and 10,150 with CpG islands. As expected, the number of DMRs is much higher compared to the monocyte/macrophage dataset. Executing the whole workflow required 30.61 hours in total with a maximum memory usage of 208.83 GB.

#### Methylome segmentation

We applied wg-blimp to a single WGBS sequencing run of H1 embryonic stem cells (ESCs) (33, 34) (SRA accession SRP072141) to demonstrate segmentation using MethylSeekR. We chose H1 embyronic stem cells to compare our integrated segmentation to the results of the original MethylSeekR authors that, among other cell types, also analyzed H1 ESCs (12). FDR cutoff was set to 5% and methylation cutoff to 50% (default values). PMDs were not considered because alpha distribution values did not suggest PMD presence in this methylome (see Supplementary Material). In total, 18,930 UMRs and 31,748 LMRs were detected.

To evaluate segmentation results, we computed each segment center’s distance to the nearest transcription start site (TSS) as reported by Ensembl (13). Figure 3 depicts separability of UMRs and LMRs with regards to TSS distances. As expected, most UMRs are in close proximity of a TSS, indicating activity in regulatory regions. Our results are in line with the original findings that also found no PMD presence and UMRs mostly overlapping promoter regions for H1 ESCs (12), despite differences in reference genomes and sequencing strategies.

**Fig. 3.**
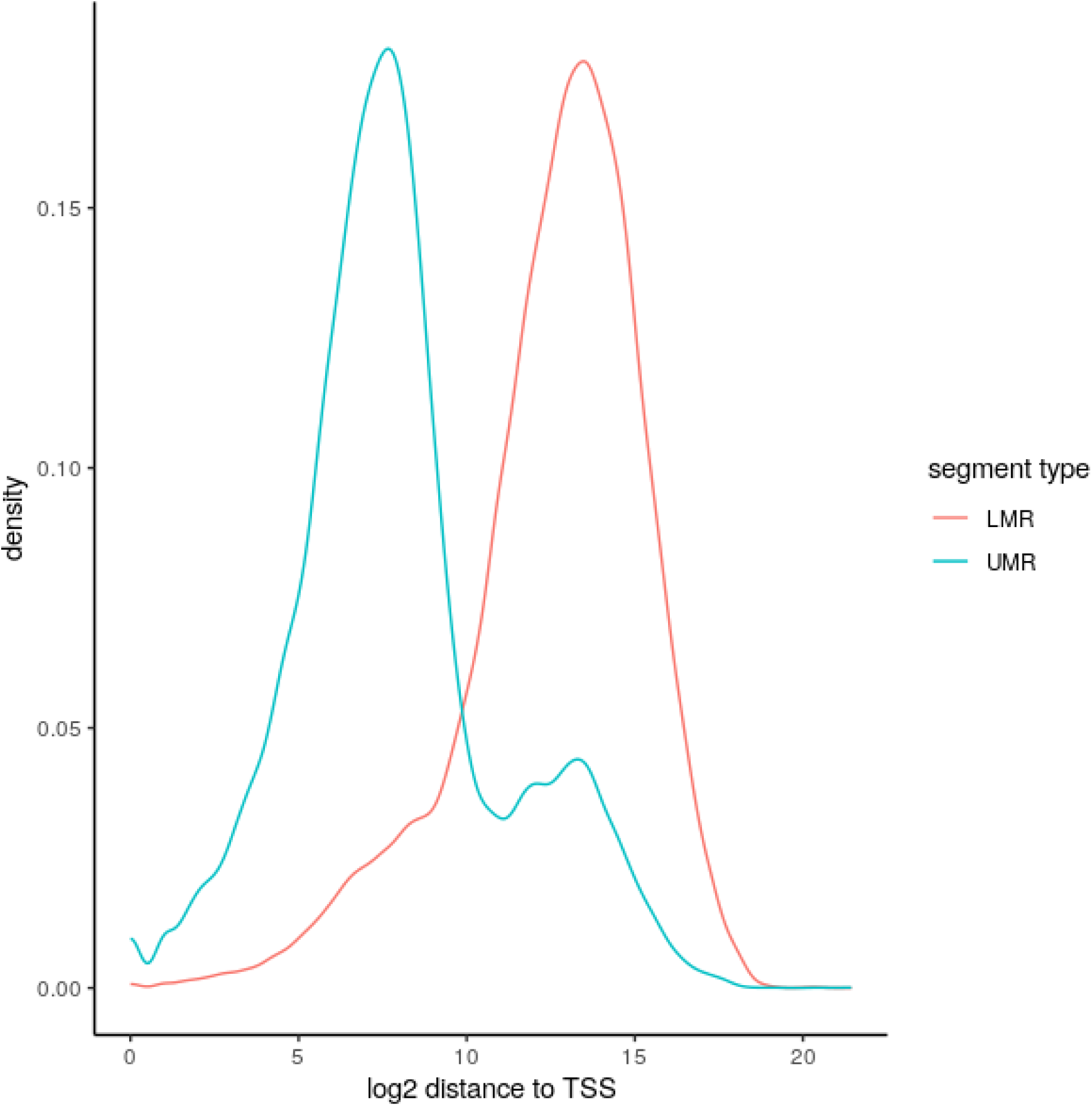
Distance from UMR/LMR centers to closest TSS for H1 ESCs. UMRs/LMRs were automatically inferred using wg-blimp’s MethylSeekR integration. UMRs and LMRs show a clear separation, with most UMRs being located in close proximity of TSSs.

Excluding reference genome indexing, executing the whole wg-blimp workflow from alignment to segmentation required 11.05 hours to complete. Alignment was the most time consuming step with a run time of 5.72 hours. Maximum memory usage of 168.76 GB was reached by MethylSeekR.

## Conclusions

wg-blimp implements a WGBS analysis workflow, improving on previous WGBS pipelines by providing simple installation and usage as well as a more extensive set of features. In addition to the analysis workflow wg-blimp includes a reactive R Shiny web interface for simplified inspection and sharing of results. wg-blimp is capable of producing coherent results, as demonstrated by analysing three publicly available datasets. We believe wg-blimp to be an apt alternative to previous WGBS analysis pipelines and hope to ease handling WGBS datasets for fellow researchers, and thus benefit the field of epigenetic research.

## Availability and requirements

**Project name:** wg-blimp.

**Project home page:** https://github.com/MarWoes/wg-blimp

**Operating system(s):** UNIX.

**Programming language:** Python, R.

**License:** AGPL-3.0

## Abbreviations

CGI: CpG island
DMR: differentially methylated region
ESC: embyronic stel cell
FDR: false-discovery rate
LMR: low-methylated region
PMD: partially methylated domain
QC: quality control
TSS: transcription start site
UMR: unmethylated region
WGBS: whole genome bisulfite sequencing

## Acknowledgements

We thank Professor Martin Dugas for support.

## Author’s contributions

MW developed the software and prepared the final version of the manuscript. EL, BH and SL reviewed analysis output and provided feedback for subsequent improvement of the software. SL further provided novel data to test the pipeline with. SR and CS tested the software and provided feedback for subsequent improvement of the software. CS provided suggestions for best-practise WGBS analysis. All authors provided feedback on the manuscript and read and approved the final version of the manuscript.

