## Supplementary Material for "wg-blimp: an end-to-end analysis pipeline for whole genome bisulfite sequencing data"

November 28, 2019

### 1 Shiny interface

We implemented a web application to ease sharing our WGBS analysis results with multiple researchers across multiple institutes and to simplify access to BAM files for inspection using the IGV. We chose the R Shiny framework for our analysis as it provides necessary functionality conveniently through R, such as loading and handling tables using the `data.table` package and creating plots with `ggplot2`.

The wg-blimp user interface consists of five separate tabs for dataset selection, quality control statistics, pipeline parameters, overview over called DMRs and UMR/LMR/PMD segmentation.

#### 1.1 Dataset selection

All WGBS datasets analysed by the wg-blimp analysis pipeline may be loaded into the Shiny user interface for sharing with other researchers. Figure S1 shows the corresponding part of the interface.

#### 1.2 QC statistics

The wg-blimp workflow performs quality control checks using FastQC, Picard, Qualimap, MultiQC and gathers methylation reports from MethylDackel. The shiny interface provides a brief overview over the gathered metrics and links to more detailed MultiQC and Qualimap reports. Links to alignment data for use with the IGV are also provided here. Figure S2 shows the statistics tab of the interface.

#### 1.3 Pipeline parameters

Results generated by bioinformatics pipelines commonly depend on the parameters in use. We provide the config file along with the DMR analysis results to enable a more compre-

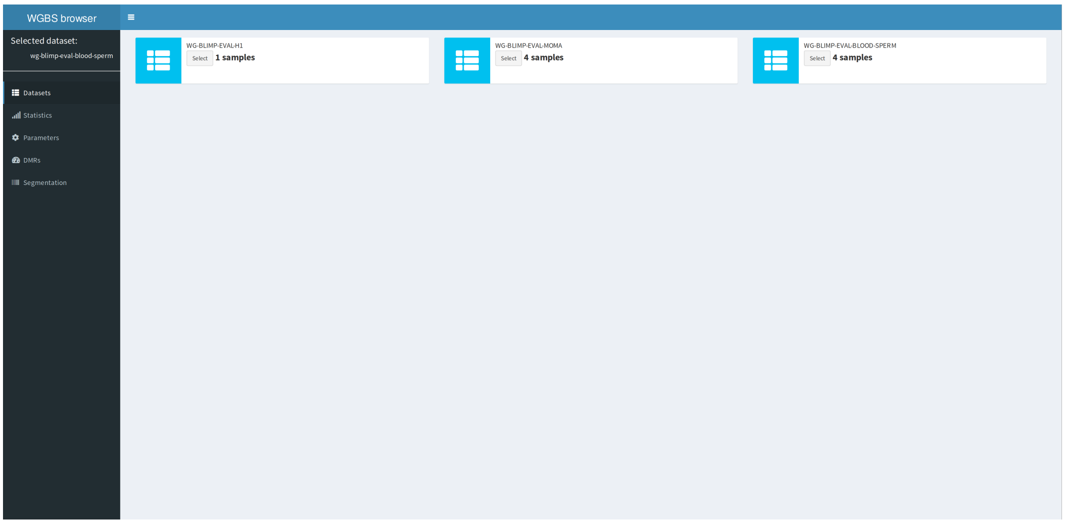


Figure S1: Shiny interface dataset selection tab.

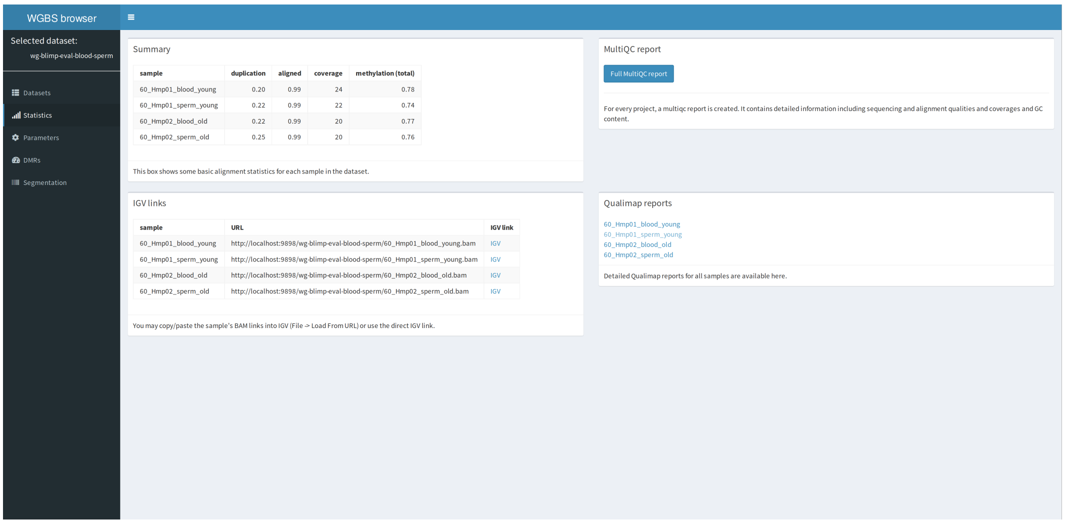


Figure S2: Shiny statistics tab.

hensive discussion of results. Figure S3 depicts the parameter overview tab.

### 1.4 DMRs

A tab to inspect DMRs retrieved by the calling pipeline is also provided. DMRs may be filtered by:

- Number of CpGs within DMR
- Absolute difference in methylation
- Length of DMR in bp

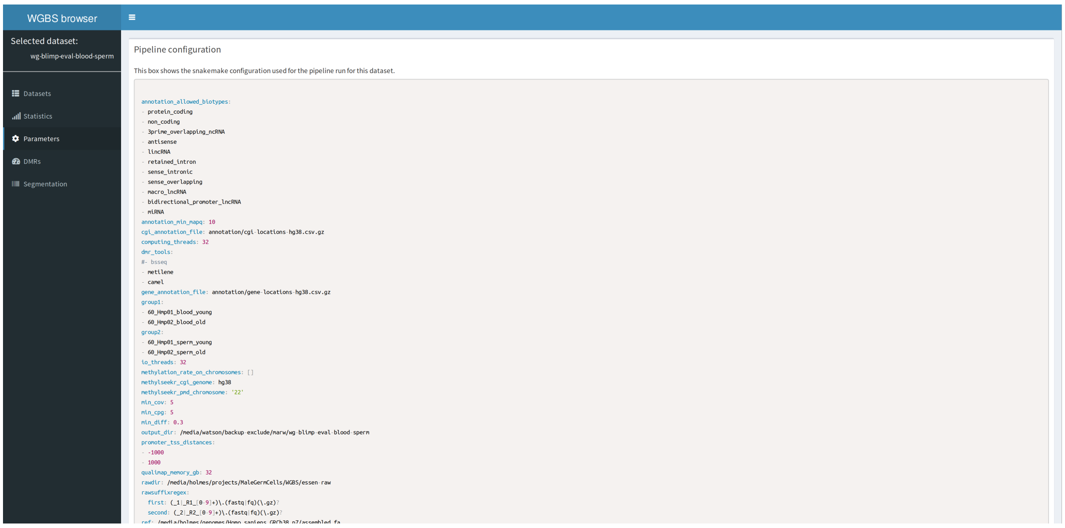

Figure S3: Shiny parameters tab.

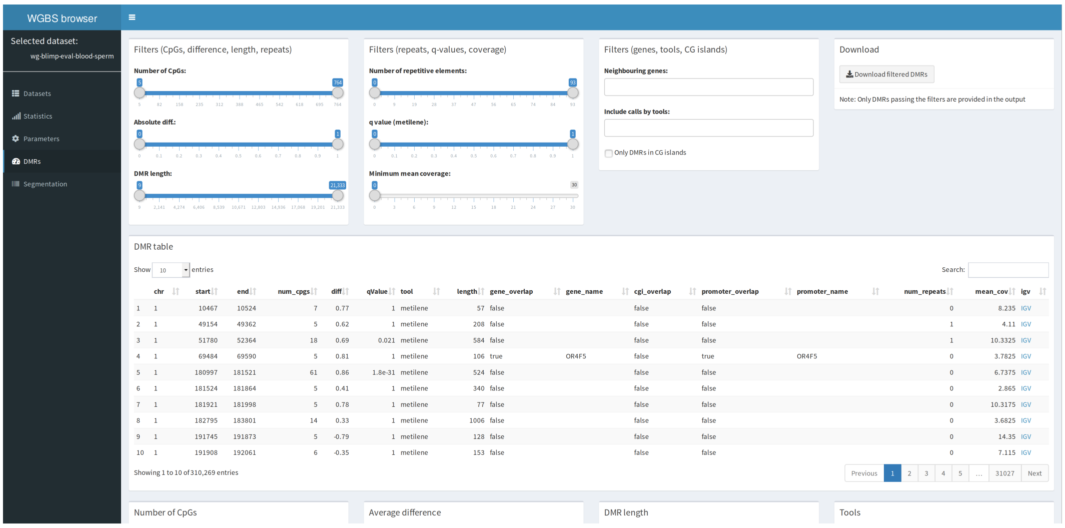

Figure S4: Shiny DMR tab.

- Number of repetitive elements within DMR
- q-value as reported by metilene
- Minimum mean coverage in DMR
- Genes (or promoters of genes) overlapped by DMRs
- Tools used for DMR calling
- Intersection with CG islands

Figure S4 gives an overview over the DMR tab. Filters are applied immediately and updated reactively through Shiny.

#### 1.5 Segmentation

wg-blimp's MethylSeekR integration preemptively computes UMRs/LMRs with and without consideration of PMDs. $\alpha$ distributions are displayed to simplify decision whether the methylome at hand contains PMDs. If the distribution appears to be bimodal, one may assume PMDs to be present. The Shiny interface also provides information on the relation of number of CpGs to median methylation in each segment, as depicted in Figure 2. Usually UMRs are longer than LMRs, which should be distinguishable from the UMR/LMR heatmap. A plot displaying information on FDR and methylation cutoffs is also provided for quality control.

### 2 Analysis of public WGBS datasets

We first applied wg-blimp to a published dataset of two isogenic pairs of human monocyte and macrophage samples. Details are described in the original publication (EGA accession EGAS00001001595) [1]. We also used wg-blimp to analyse a blood and sperm dataset to find differences in methylation between sperm and somatic cells. The dataset is also publicly available (ENA accession PRJEB28044). For this dataset pooled DNA from sperm and blood DNA (n=6 for each) was extracted from healthy young (age 18-24) and old (age 61-71) men, yielding four WGBS samples [2]. For both datasets 1% Lambda phage DNA was added for later quality control. Sequencing was performed on Illumina HiSeq 2500. Protocols for the H1 ESC sample are described in the original publication (NCBI SRA accession SRR3274347) [3].

#### 2.1 Quality control

Table S1 shows the QC parameters as displayed in wg-blimp's shiny interface for each of the datasets.

Coverage profiles are $\geq 19\times$ and the mean methylation for lambda phage indicates successful bisulfite conversion for the monocyte/macrophage and blood/sperm datasets. Please note that lambda phage conversion rates are lacking for the H1 ESC sample as reads assigned to the lambda genome were discarded during upload to read repositories and thus not included in our computation. Since no QC parameter suggests errors, all samples were used for further analysis.

#### 2.2 DMRs

The DMR algorithms used are expected to yield low numbers of calls for the monocyte/-macrophage dataset and high numbers for the blood/sperm comparison. Filtering may be performed in a trivial fashion through wg-blimp's user interface.

| dataset | sample | duplication | aligned | coverage | conversion | methylation |
|---|---|---|---|---|---|---|
| monocyte / macrophage | 43_Hm03_BlMa | 0.23 | 1.00 | 22 | 0.98 | 0.80 |
| monocyte / macrophage | 43_Hm03_BlMo | 0.19 | 1.00 | 21 | 0.99 | 0.80 |
| monocyte / macrophage | 43_Hm05_BlMa | 0.23 | 1.00 | 33 | 0.99 | 0.81 |
| monocyte / macrophage | 43_Hm05_BlMo | 0.22 | 1.00 | 33 | 0.99 | 0.81 |
| blood / sperm | blood (young) | 0.20 | 0.99 | 24 | 0.99 | 0.78 |
| blood / sperm | sperm (young) | 0.22 | 0.99 | 22 | 0.99 | 0.74 |
| blood / sperm | blood (old) | 0.22 | 0.99 | 20 | 0.99 | 0.77 |
| blood / sperm | sperm (old) | 0.25 | 0.99 | 20 | 1.00 | 0.76 |
| H1 ESC | SRR3274347 | 0.05 | 1.00 | 19 | - | 0.76 |

Table S1: Quality control summary of analysed public datasets

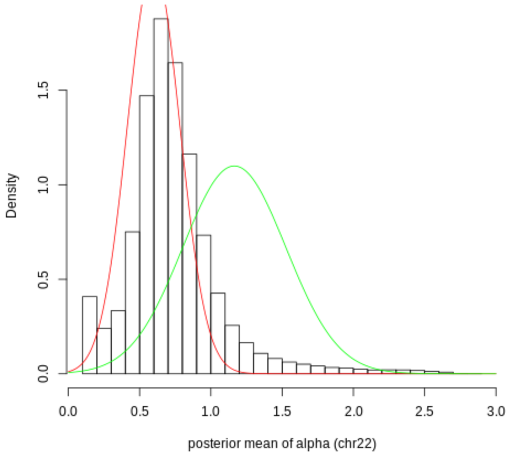


Figure S5: Distribution of $\alpha$ values for chromosome 22 of H1 ESC sample

### 2.3 Segmentation

We focus on segmenting the H1 ESC methylome because of its previous analysis by the authors of the original MethylSeekR publication [4]. $\alpha$ distributions are similar for different chromosomes, so MethylSeekR infers the distribution for only a single chromosome. $\alpha$ smaller than 1 represents a polarized distribution favoring low and high methylation. $\alpha$ equal or greater than 1 suggests a uniform distribution. If an $\alpha$ distribution has a bimodal shape, we assume presence of PMDs. Since the $\alpha$ distribution appears to be unimodal with only few $\alpha$ values $> 1$, we do not consider PMDs for the H1 ESC sample, as depicted in Figure S5. This is consistent with the analysis of the original MethylSeekR authors where H1 cells showed a similar $\alpha$ distribution whereas, among other, fetal lung fibroblasts showed a clearly distinguishable bimodal $\alpha$ distribution [4]. Figure S6 shows intersection of LMRs/UMRs with exons and promoters. We here set promoters to the ranges around transcription start sites (TSSs): $[TSS - 1000, TSS + 1000]$. As expected, while 82.32% of UMRs are intersecting promoters, only 12.29% of LMRs show promoter intersection. This is also coherent with the original MethylSeekR findings [4].

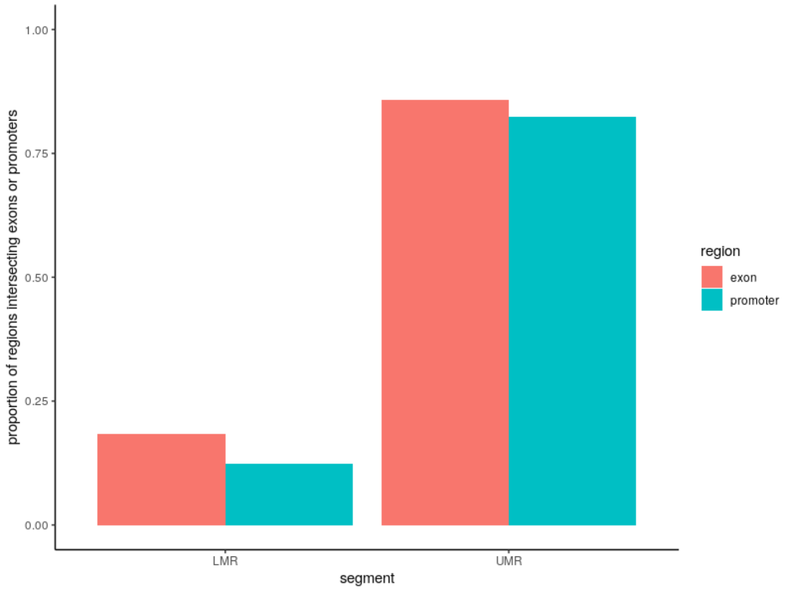


Figure S6: Intersection of LMRs/UMRs with exons and promoters

### 2.4 Computational requirements

wg-blimp logs run times and memory requirements for each step. The resulting plots are displayed in Figures S7 to S12.

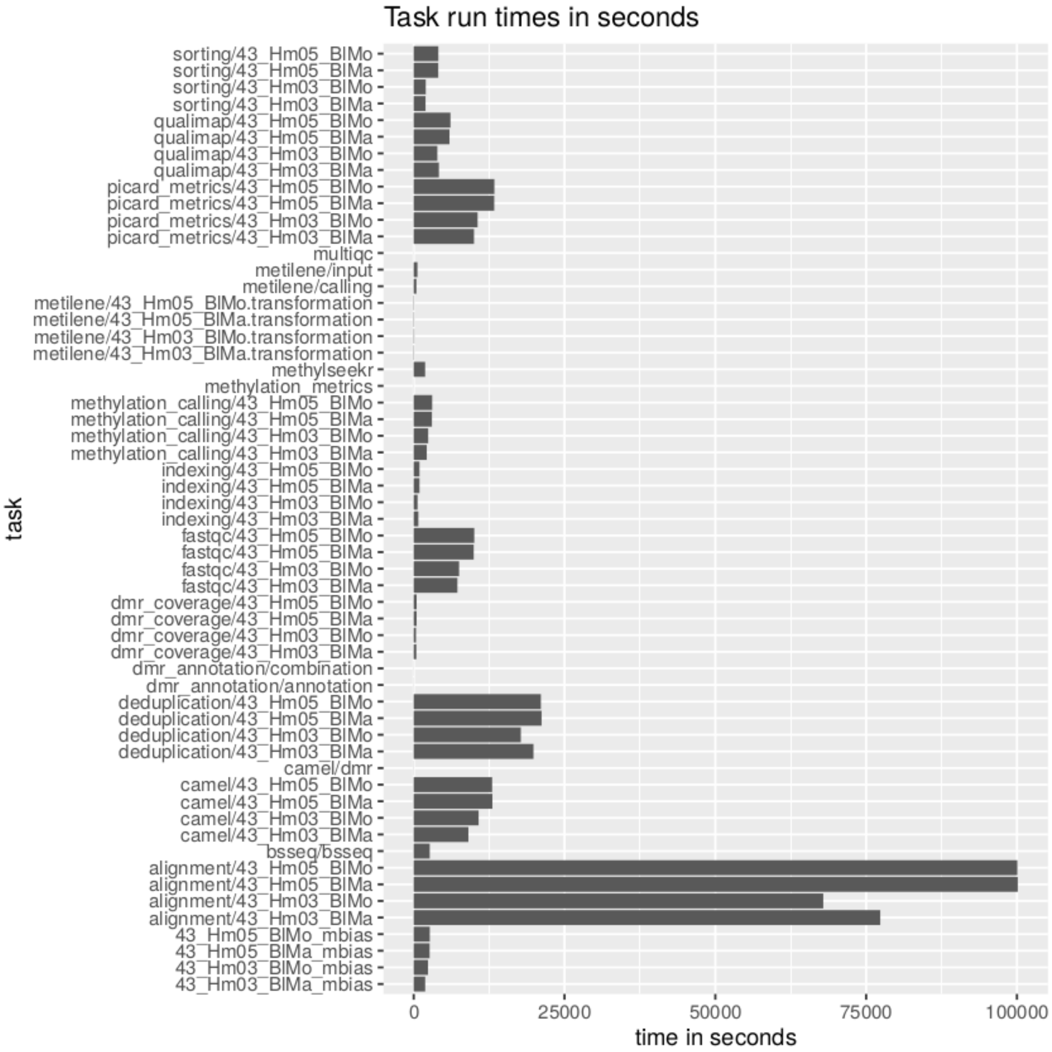


Figure S7: Run times for monocyte/macrophage dataset

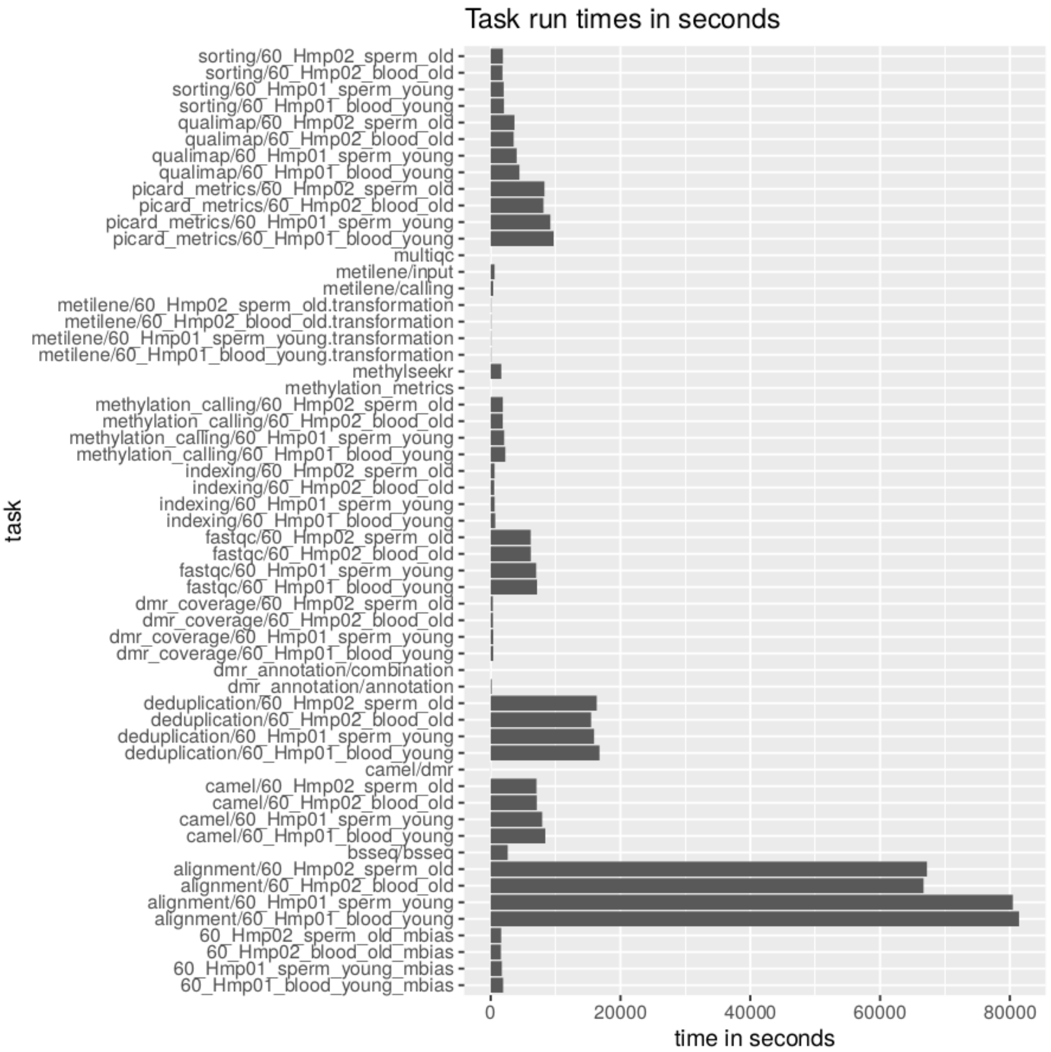


Figure S8: Run times for blood/sperm dataset

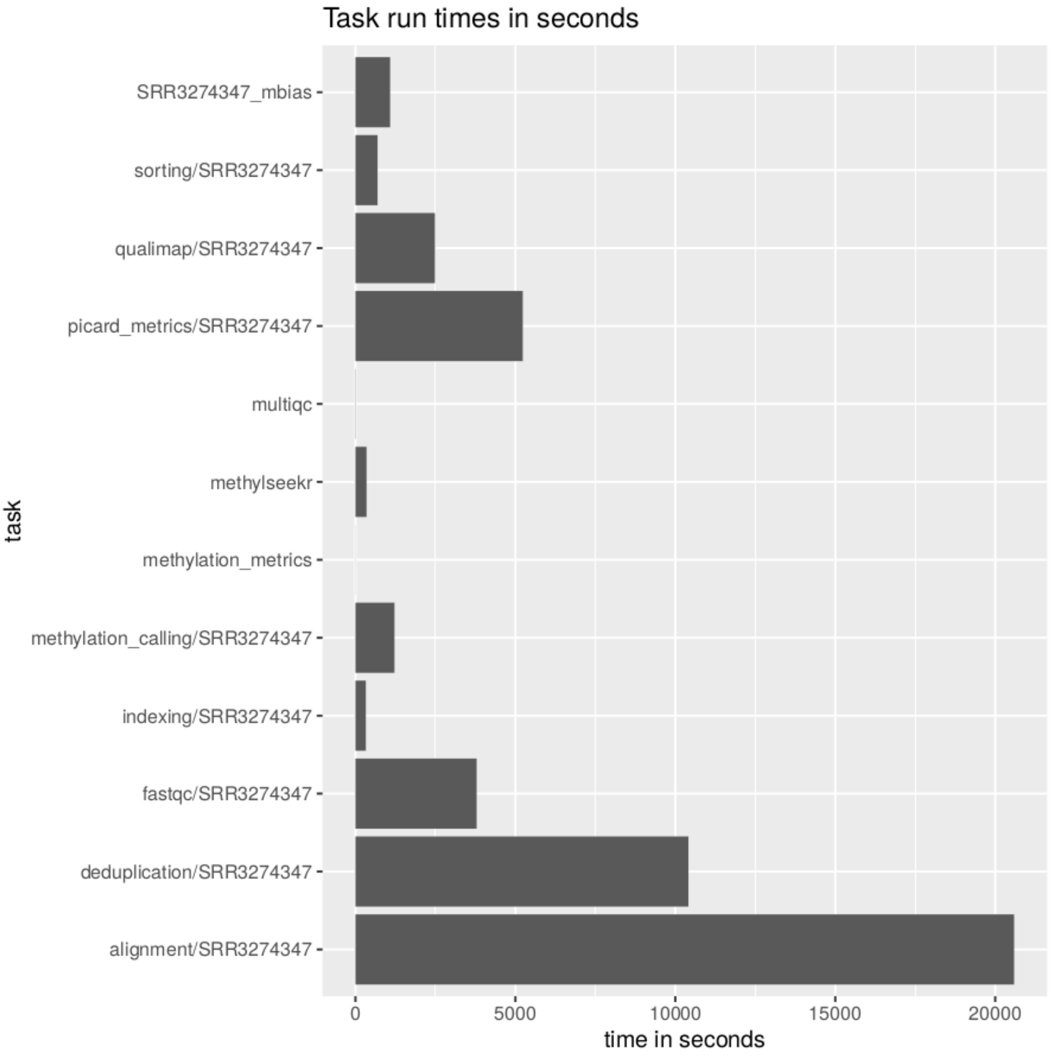


Figure S9: Run times for H1 ESC sample

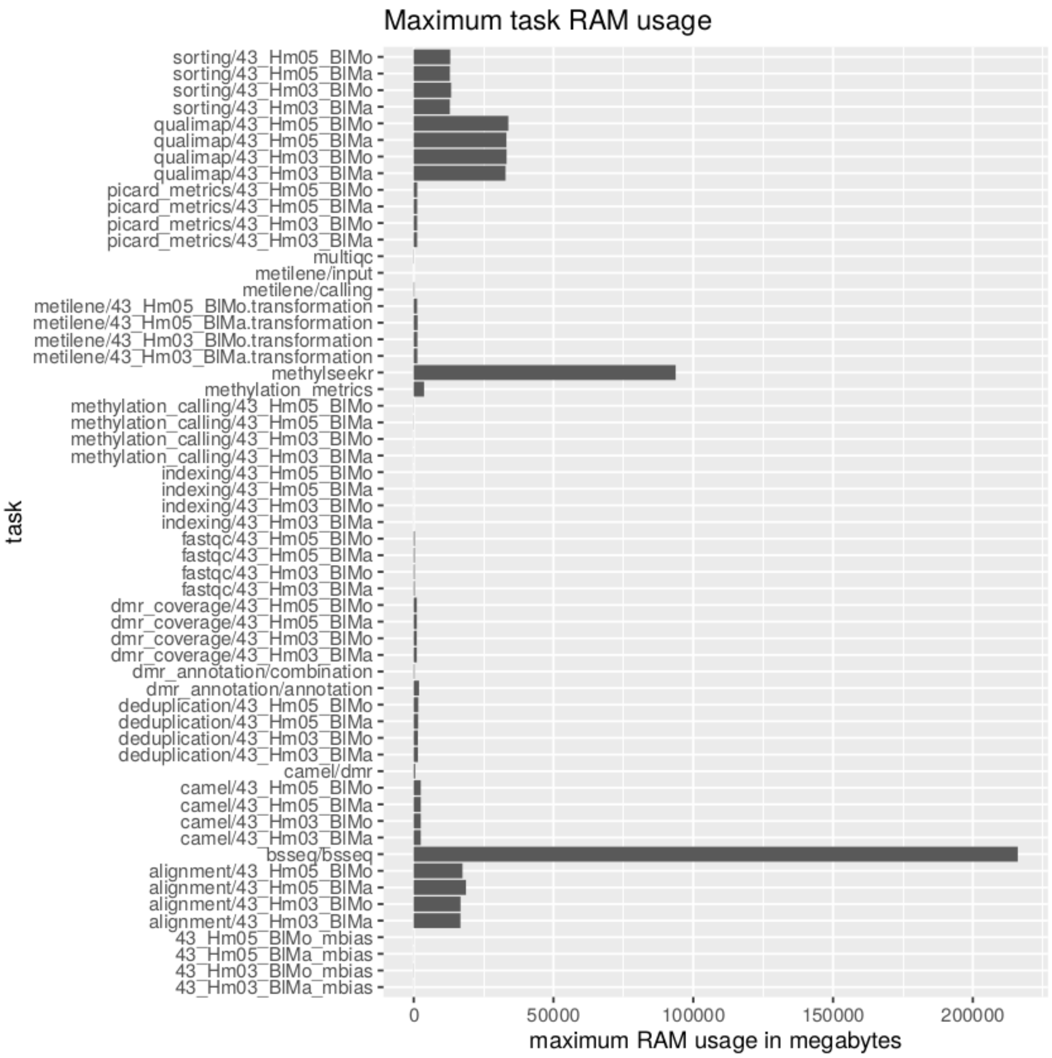


Figure S10: Memory usage for monocyte/macrophage dataset

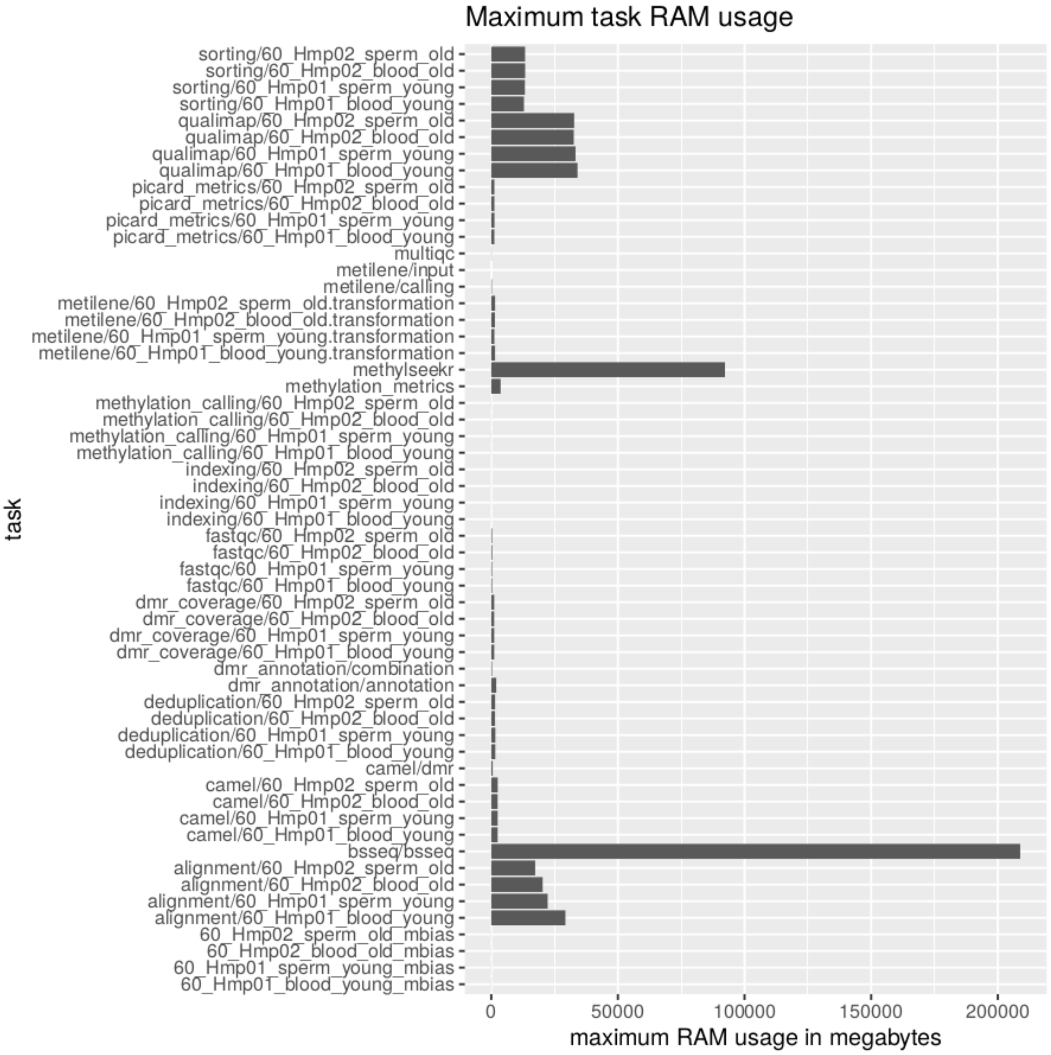


Figure S11: Memory usage for blood/sperm dataset

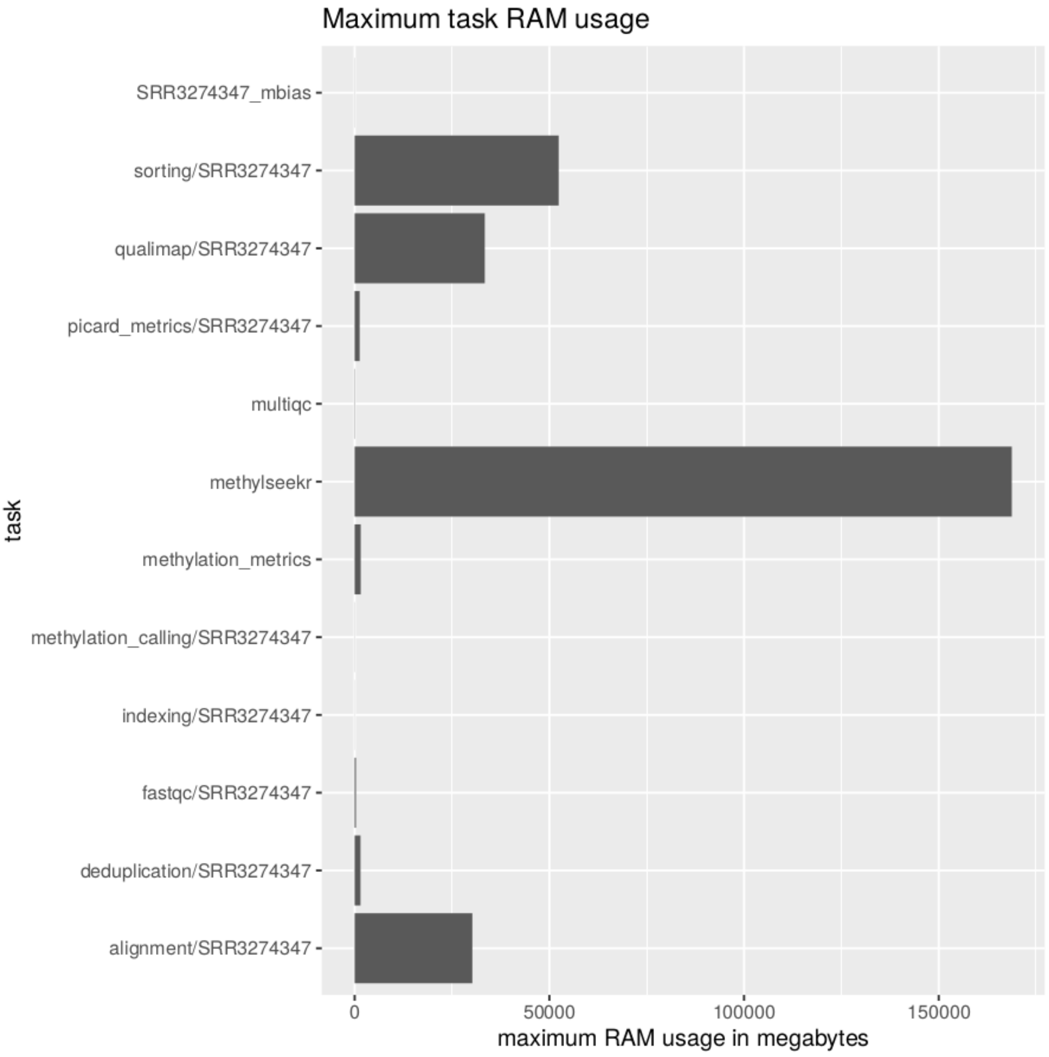


Figure S12: Memory usage for H1 ESC sample

### 2.5 Parameters

This section shows the configuration files and environment used for the analyses.

#### 2.5.1 Monocyte/Macrophage configuration file

```
annotation_allowed_biotypes:
- protein_coding
- non_coding
- 3prime_overlapping_ncRNA
- antisense
- lincRNA
- retained_intron
- sense_intronic
- sense_overlapping
- macro_lncRNA
- bidirectional_promoter_lncRNA
- miRNA
annotation_min_mapq: 10
bsseq_local_correct: false
cgi_annotation_file: annotation/cgi-locations-hg38.csv.gz
computing_threads: 16
dmr_tools:
- bsseq
- camel
- metilene
gene_annotation_file: annotation/gene-locations-hg38.csv.gz
group1:
- 43_Hm03_BlMa
- 43_Hm05_BlMa
group2:
- 43_Hm03_BlMo
- 43_Hm05_BlMo
io_threads: 16
methylation_rate_on_chromosomes: []
methylseekr_cgi_genome: hg38
methylseekr_fdr_cutoff: 5
methylseekr_methylation_cutoff: 0.5
methylseekr_pmd_chromosome: '22'
min_cov: 5
min_cpg: 4
min_diff: 0.3
output_dir: /media/watson/backup-exclude/marw/wg-blimp-eval-moma
promoter_tss_distances:
- -1000
- 1000
qualimap_memory_gb: 32
rawdir: /media/holmes/projects/MaleGermCells/WGBS/essen-raw
rawsuffixregex:
  first: (_1|_R1_[0-9]+)\.(fastq|fq)(\.gz)?
  second: (_2|_R2_[0-9]+)\.(fastq|fq)(\.gz)?
ref: /media/holmes/genomes/Homo_sapiens.GRCh38.p7/assembled.fa
repeat_masker_annotation_file: annotation/repeat-masker-hg38.csv.gz
repeat_masker_links:
  annotation/repeat-masker-hg19.csv.gz: https://uni-muenster.sciebo.de/s/tFPOT7weAc5eGLG/download
  annotation/repeat-masker-hg38.csv.gz: https://uni-muenster.sciebo.de/s/LwMik3kKY83oVT0/download
samples:
- 43_Hm03_BlMa
- 43_Hm05_BlMa
- 43_Hm03_BlMo
- 43_Hm05_BlMo
target_files:
- dmr/annotated-dmrs.csv
- qc/multiqc_report.html
- qc/methylation_metrics.csv
- segmentation/umr-lmr-all.csv
transcript_start_site_file: annotation/transcription-start-sites-hg38.csv.gz
```

#### 2.5.2 Blood/Sperm configuration file

```
annotation_allowed_biotypes:
- protein_coding
- non_coding
- 3prime_overlapping_ncRNA
- antisense
- lincRNA
- retained_intron
- sense_intronic
- sense_overlapping
- macro_lncRNA
```

```
- bidirectional_promoter_lncRNA
- miRNA
annotation_min_mapq: 10
bsseq_local_correct: false
cgi_annotation_file: annotation/cgi-locations-hg38.csv.gz
computing_threads: 16
dmr_tools:
- bsseq
- camel
- metilene
gene_annotation_file: annotation/gene-locations-hg38.csv.gz
group1:
- 60_Hmp01_blood_young
- 60_Hmp02_blood_old
group2:
- 60_Hmp01_sperm_young
- 60_Hmp02_sperm_old
io_threads: 16
methylation_rate_on_chromosomes: ['LAMBDA']
methylseekr_cgi_genome: hg38
methylseekr_fdr_cutoff: 5
methylseekr_methylation_cutoff: 0.5
methylseekr_pmd_chromosome: '22'
min_cov: 5
min_cpg: 4
min_diff: 0.3
output_dir: /media/watson/backup-exclude/marw/wg-blimp-eval-blood-sperm
promoter_tss_distances:
- -1000
- 1000
qualimap_memory_gb: 32
rawdir: /media/holmes/projects/MaleGermCells/WGBS/essen-raw
rawsuffixregex:
  first: (_1|_R1_[0-9]+)\.(fastq|fq)(\.gz)?
  second: (_2|_R2_[0-9]+)\.(fastq|fq)(\.gz)?
ref: /media/holmes/genomes/Homo_sapiens.GRCh38.p7/assembled.fa
repeat_masker_annotation_file: annotation/repeat-masker-hg38.csv.gz
repeat_masker_links:
  annotation/repeat-masker-hg19.csv.gz: https://uni-muenster.sciebo.de/s/tFPOT7weAc5eGLG/download
  annotation/repeat-masker-hg38.csv.gz: https://uni-muenster.sciebo.de/s/LwMik3kKY83oVT0/download
samples:
- 60_Hmp01_blood_young
- 60_Hmp02_blood_old
- 60_Hmp01_sperm_young
- 60_Hmp02_sperm_old
target_files:
- dmr/annotated-dmrs.csv
- qc/multiqc_report.html
- qc/methylation_metrics.csv
- segmentation/umr-lmr-all.csv
transcript_start_site_file: annotation/transcription-start-sites-hg38.csv.gz
```

#### 2.5.3 H1 ESC configuration file

```
annotation_allowed_biotypes:
- protein_coding
- non_coding
- 3prime_overlapping_ncRNA
- antisense
- lincRNA
- retained_intron
- sense_intronic
- sense_overlapping
- macro_lncRNA
- bidirectional_promoter_lncRNA
- miRNA
annotation_min_mapq: 10
bsseq_local_correct: false
cgi_annotation_file: annotation/cgi-locations-hg38.csv.gz
computing_threads: 64
dmr_tools:
- bsseq
gene_annotation_file: annotation/gene-locations-hg38.csv.gz
group1:
- SRR3274347
group2:
- SRR3274347
io_threads: 64
methylation_rate_on_chromosomes: []
methylseekr_cgi_genome: hg38
methylseekr_fdr_cutoff: 5
methylseekr_methylation_cutoff: 0.5
methylseekr_pmd_chromosome: '22'
min_cov: 5
```

```
min_cpg: 5
min_diff: 0.3
output_dir: /media/holmes/backup-exclude/wg-blimp-eval-h1
promoter_tss_distances:
- -1000
- 1000
qualimap_memory_gb: 32
rawdir: /media/holmes/backup-exclude/wg-blimp-eval-h1/fastq
rawsuffixregex:
  first: (_1|_R1_[0-9]+)\.(fastq|fq)(\.gz)?
  second: (_2|_R2_[0-9]+)\.(fastq|fq)(\.gz)?
ref: /media/holmes/genomes/Homo_sapiens.GRCh38.p7/assembled.fa
repeat_masker_annotation_file: annotation/repeat-masker-hg38.csv.gz
repeat_masker_links:
  annotation/repeat-masker-hg19.csv.gz: https://uni-muenster.sciebo.de/s/tFPOT7weAc5eGLG/download
  annotation/repeat-masker-hg38.csv.gz: https://uni-muenster.sciebo.de/s/LwMik3kKY83oVT0/download
samples:
- SRR3274347
target_files:
- qc/multiqc_report.html
- qc/methylation_metrics.csv
- segmentation/umr-lmr-all.csv
transcript_start_site_file: annotation/transcription-start-sites-hg38.csv.gz
```

#### 2.5.4 wg-blimp conda environment

```
Name                          Version                Build  Channel
_libgcc_mutex                 0.1                     main
_r-mutex                      1.0.1            anacondar_1  conda-forge
aioeasywebdav                 2.4.0              py36_1000  conda-forge
aiohttp                       3.5.4       py36h14c3975_0    conda-forge
alsa-lib                      1.1.5          h516909a_1001  conda-forge
appdirs                       1.4.3                   py_1  conda-forge
asn1crypto                    0.24.0             py36_1003  conda-forge
async-timeout                 3.0.1                py_1000  conda-forge
attrs                         19.1.0                  py_0  conda-forge
bcrypt                        3.1.6         py36h516909a_1  conda-forge
bedtools                      2.28.0            hdf88d34_0  bioconda
binutils_impl_linux-64        2.31.1            h6176602_1
binutils_linux-64             2.31.1            h6176602_8
bioconductor-annotate         1.62.0                 r36_1  bioconda
bioconductor-annotationdbi 1.46.0                 r36_1  bioconda
bioconductor-beachmat         2.0.0         r36he1b5a44_1  bioconda
bioconductor-biobase          2.44.0        r36h516909a_1  bioconda
bioconductor-biocgenerics 0.30.0                 r36_1  bioconda
bioconductor-biocparallel 1.18.0         r36he1b5a44_1  bioconda
bioconductor-biostrings       2.52.0        r36h516909a_1  bioconda
bioconductor-bsgenome         1.52.0                 r36_1  bioconda
bioconductor-bsseq            1.20.0        r36he1b5a44_1  bioconda
bioconductor-delayedarray 0.10.0         r36h516909a_1  bioconda
bioconductor-delayedmatrixstats 1.6.0                 r36_1  bioconda
bioconductor-geneplotter      1.62.0                 r36_1  bioconda
bioconductor-genomeinfodb 1.20.0                 r36_1  bioconda
bioconductor-genomeinfodbdata 1.2.1                 r36_1  bioconda
bioconductor-genomicalignments 1.20.1        r36h516909a_0  bioconda
bioconductor-genomicranges 1.36.0        r36h516909a_1  bioconda
bioconductor-hdf5array        1.12.1        r36h516909a_0  bioconda
bioconductor-iranges          2.18.1        r36h516909a_0  bioconda
bioconductor-limma            3.40.2        r36h516909a_0  bioconda
bioconductor-methylseekr      1.24.0                 r36_1  bioconda
bioconductor-noiseq           2.28.0                 r36_1  bioconda
bioconductor-rhdf5            2.28.0        r36he1b5a44_1  bioconda
bioconductor-rhdf5lib         1.6.0         r36h516909a_1  bioconda
bioconductor-rhtslib          1.16.1        r36hbcae180_1  bioconda
bioconductor-rsamtools        2.0.0         r36he1b5a44_1  bioconda
bioconductor-rtracklayer      1.44.2        r36h516909a_1  bioconda
bioconductor-s4vectors        0.22.0        r36h516909a_1  bioconda
bioconductor-summarizedexperiment 1.14.0                 r36_1  bioconda
bioconductor-xvector          0.24.0        r36h516909a_1  bioconda
bioconductor-zlibbioc         1.30.0        r36h516909a_1  bioconda
boto3                         1.9.211                 py_0  conda-forge
botocore                      1.12.211                py_0  conda-forge
bwa                           0.7.17            hed695b0_6  bioconda
bwameth                       0.2.0                py36_1  bioconda
bwidget                       1.9.11                     0  conda-forge
bzip2                         1.0.8             h516909a_0  conda-forge
ca-certificates               2019.6.16         hecc5488_0  conda-forge
cachetools                    2.1.0                   py_0  conda-forge
cairo                         1.16.0         h18b612c_1001  conda-forge
certifi                       2019.6.16            py36_1  conda-forge
cffi                          1.12.3        py36h8022711_0  conda-forge
chardet                       3.0.4              py36_1003  conda-forge
click                         7.0                     py_0  conda-forge
colormath                     3.0.0                   py_2  conda-forge
```

| | | | |
|---|---|---|---|
| configargparse | 0.13.0 | py_1 | conda-forge |
| cryptography | 2.7 | py36h72c5cf5_0 | conda-forge |
| curl | 7.65.3 | hf8cf82a_0 | conda-forge |
| cycler | 0.10.0 | py_1 | conda-forge |
| datrie | 0.8 | py36h516909a_0 | conda-forge |
| dbus | 1.13.6 | he372182_0 | conda-forge |
| ddt | 1.2.1 | pypi_0 | pypi |
| decorator | 4.4.0 | py_0 | conda-forge |
| docutils | 0.15.2 | py36_0 | conda-forge |
| dropbox | 9.4.0 | py_0 | conda-forge |
| expat | 2.2.5 | he1b5a44_1003 | conda-forge |
| fastqc | 0.11.8 | 1 | bioconda |
| filechunkio | 1.8 | py_2 | conda-forge |
| font-ttf-dejavu-sans-mono | 2.37 | h6964260_0 | |
| fontconfig | 2.13.1 | he4413a7_1000 | conda-forge |
| freetype | 2.10.0 | he983fc9_1 | conda-forge |
| ftputil | 3.4 | py_0 | conda-forge |
| future | 0.17.1 | py36_1000 | conda-forge |
| gcc_impl_linux-64 | 7.3.0 | habb00fd_1 | conda-forge |
| gcc_linux-64 | 7.3.0 | h553295d_8 | conda-forge |
| gettext | 0.19.8.1 | hc5be6a0_1002 | conda-forge |
| gfortran_impl_linux-64 | 7.3.0 | hdf63c60_1 | |
| gfortran_linux-64 | 7.3.0 | h553295d_8 | |
| giflib | 5.1.7 | h516909a_1 | conda-forge |
| gitdb2 | 2.0.5 | py_0 | conda-forge |
| gitpython | 3.0.1 | py_0 | conda-forge |
| glib | 2.58.3 | h6f030ca_1002 | conda-forge |
| google-api-core | 1.14.2 | py36_0 | conda-forge |
| google-auth | 1.6.3 | py_0 | conda-forge |
| google-cloud-core | 1.0.3 | py_0 | conda-forge |
| google-cloud-storage | 1.18.0 | py_0 | conda-forge |
| google-resumable-media | 0.3.2 | py_0 | conda-forge |
| googleapis-common-protos | 1.6.0 | py36_0 | conda-forge |
| graphite2 | 1.3.13 | hf484d3e_1000 | conda-forge |
| graphviz | 2.38.0 | hf68f40c_1011 | conda-forge |
| gsl | 2.5 | h294904e_0 | conda-forge |
| gst-plugins-base | 1.14.5 | h0935bb2_0 | conda-forge |
| gstreamer | 1.14.5 | h36ae1b5_0 | conda-forge |
| gxx_impl_linux-64 | 7.3.0 | hdf63c60_1 | conda-forge |
| gxx_linux-64 | 7.3.0 | h553295d_8 | conda-forge |
| h5py | 2.9.0 | nompi_py36h513d04c_1104 | conda-forge |
| harfbuzz | 2.4.0 | h37c48d4_1 | conda-forge |
| hdf5 | 1.10.5 | nompi_h3c11f04_1102 | conda-forge |
| htslib | 1.9 | ha228f0b_7 | bioconda |
| icu | 58.2 | hf484d3e_1000 | conda-forge |
| idna | 2.8 | py36_1000 | conda-forge |
| idna_ssl | 1.1.0 | py36_1000 | conda-forge |
| jinja2 | 2.10.1 | py_0 | conda-forge |
| jmespath | 0.9.4 | py_0 | conda-forge |
| jpeg | 9c | h14c3975_1001 | conda-forge |
| jsonschema | 3.0.2 | py36_0 | conda-forge |
| kiwisolver | 1.1.0 | py36hc9558a2_0 | conda-forge |
| krb5 | 1.16.3 | h05b26f9_1001 | conda-forge |
| lcms2 | 2.9 | h2e4bb80_0 | conda-forge |
| libblas | 3.8.0 | 12_openblas | conda-forge |
| libcblas | 3.8.0 | 12_openblas | conda-forge |
| libcurl | 7.65.3 | hda55be3_0 | conda-forge |
| libdeflate | 1.0 | h14c3975_1 | bioconda |
| libedit | 3.1.20170329 | hf8c457e_1001 | conda-forge |
| libffi | 3.2.1 | he1b5a44_1006 | conda-forge |
| libgcc-ng | 9.1.0 | hdf63c60_0 | |
| libgfortran-ng | 7.3.0 | hdf63c60_0 | |
| libiconv | 1.15 | h516909a_1005 | conda-forge |
| liblapack | 3.8.0 | 12_openblas | conda-forge |
| libopenblas | 0.3.7 | h6e990d7_1 | conda-forge |
| libpng | 1.6.37 | hed695b0_0 | conda-forge |
| libprotobuf | 3.9.1 | h8b12597_0 | conda-forge |
| libssh2 | 1.8.2 | h22169c7_2 | conda-forge |
| libstdcxx-ng | 9.1.0 | hdf63c60_0 | |
| libtiff | 4.0.10 | h57b8799_1003 | conda-forge |
| libtool | 2.4.6 | h14c3975_1002 | conda-forge |
| libuuid | 2.32.1 | h14c3975_1000 | conda-forge |
| libxcb | 1.13 | h14c3975_1002 | conda-forge |
| libxml2 | 2.9.9 | h13577e0_2 | conda-forge |
| lz4-c | 1.8.3 | he1b5a44_1001 | conda-forge |
| lzstring | 1.0.4 | py_1001 | conda-forge |
| make | 4.2.1 | h14c3975_2004 | conda-forge |
| markdown | 3.1.1 | py_0 | conda-forge |
| markupsafe | 1.1.1 | py36h14c3975_0 | conda-forge |
| matplotlib | 3.1.1 | py36_0 | conda-forge |
| matplotlib-base | 3.1.1 | py36hfd891ef_0 | conda-forge |
| methyldackel | 0.4.0 | hc0aa232_0 | bioconda |
| metilene | 0.2.6 | h14c3975_2 | bioconda |
| mosdepth | 0.2.5 | hb763d49_0 | bioconda |
| multidict | 4.5.2 | py36h14c3975_1000 | conda-forge |

| | | | |
|---|---|---|---|
| multiqc | 1.7 | py_4 | bioconda |
| ncurses | 6.1 | hf484d3e_1002 | conda-forge |
| networkx | 2.3 | py_0 | conda-forge |
| numpy | 1.17.0 | py36h95a1406_0 | conda-forge |
| openjdk | 11.0.1 | h46a85a0_1017 | conda-forge |
| openssl | 1.1.1c | h516909a_0 | conda-forge |
| pandas | 0.25.0 | py36hb3f55d8_0 | conda-forge |
| pango | 1.40.14 | he7ab937_1005 | conda-forge |
| paramiko | 2.6.0 | py36_0 | conda-forge |
| pcre | 8.41 | hf484d3e_1003 | conda-forge |
| perl | 5.26.2 | h516909a_1006 | conda-forge |
| picard | 2.20.5 | 0 | bioconda |
| pip | 19.2.2 | py36_0 | conda-forge |
| pixman | 0.38.0 | h516909a_1003 | conda-forge |
| prettytable | 0.7.2 | py_3 | conda-forge |
| protobuf | 3.9.1 | py36he1b5a44_0 | conda-forge |
| psutil | 5.6.3 | py36h516909a_0 | conda-forge |
| pthread-stubs | 0.4 | h14c3975_1001 | conda-forge |
| pyasn1 | 0.4.6 | py_0 | conda-forge |
| pyasn1-modules | 0.2.6 | py_0 | conda-forge |
| pycparser | 2.19 | py36_1 | conda-forge |
| pygments | 2.4.2 | py_0 | conda-forge |
| pygraphviz | 1.5 | py36h516909a_1001 | conda-forge |
| pynacl | 1.3.0 | py36h14c3975_1000 | conda-forge |
| pyopenssl | 19.0.0 | py36_0 | conda-forge |
| pyparsing | 2.4.2 | py_0 | conda-forge |
| pyqt | 5.9.2 | py36hcca6a23_2 | conda-forge |
| pyrsistent | 0.15.4 | py36h516909a_0 | conda-forge |
| pysam | 0.15.3 | py36hda2845c_1 | bioconda |
| pysftp | 0.2.9 | py_1 | conda-forge |
| pysocks | 1.7.0 | py36_0 | conda-forge |
| python | 3.6.7 | h357f687_1005 | conda-forge |
| python-dateutil | 2.8.0 | py_0 | conda-forge |
| python-irodsclient | 0.7.0 | py_0 | conda-forge |
| pytz | 2019.2 | py_0 | conda-forge |
| pyyaml | 5.1.2 | py36h516909a_0 | conda-forge |
| qt | 5.9.7 | h52cfd70_2 | conda-forge |
| qualimap | 2.2.2c | 1 | bioconda |
| r-assertthat | 0.2.1 | r36h6115d3f_1 | conda-forge |
| r-backports | 1.1.4 | r36hcdcec82_1 | conda-forge |
| r-base | 3.6.1 | h8900bf8_2 | conda-forge |
| r-bh | 1.69.0_1 | r36h6115d3f_1 | conda-forge |
| r-bit | 1.1_14 | r36hcdcec82_1 | conda-forge |
| r-bit64 | 0.9_7 | r36hcdcec82_1001 | conda-forge |
| r-bitops | 1.0_6 | r36hcdcec82_1003 | conda-forge |
| r-blob | 1.2.0 | r36_1 | conda-forge |
| r-cli | 1.1.0 | r36h6115d3f_1 | conda-forge |
| r-colorspace | 1.4_1 | r36hcdcec82_1 | conda-forge |
| r-crayon | 1.3.4 | r36h6115d3f_1002 | conda-forge |
| r-crosstalk | 1.0.0 | r36h6115d3f_1002 | conda-forge |
| r-data.table | 1.12.2 | r36hcdcec82_1 | conda-forge |
| r-dbi | 1.0.0 | r36h6115d3f_1002 | conda-forge |
| r-digest | 0.6.20 | r36h0357c0b_1 | conda-forge |
| r-dt | 0.8 | r36h6115d3f_0 | conda-forge |
| r-ellipsis | 0.2.0.1 | r36hcdcec82_1 | conda-forge |
| r-fansi | 0.4.0 | r36hcdcec82_1001 | conda-forge |
| r-formatr | 1.7 | r36h6115d3f_1 | conda-forge |
| r-futile.logger | 1.4.3 | r36h6115d3f_1002 | conda-forge |
| r-futile.options | 1.0.1 | r36h6115d3f_1001 | conda-forge |
| r-getopt | 1.20.3 | r36_1 | conda-forge |
| r-ggplot2 | 3.2.1 | r36h6115d3f_0 | conda-forge |
| r-glue | 1.3.1 | r36hcdcec82_1 | conda-forge |
| r-gridextra | 2.3 | r36h6115d3f_1002 | conda-forge |
| r-gtable | 0.3.0 | r36h6115d3f_2 | conda-forge |
| r-gtools | 3.8.1 | r36hcdcec82_1003 | conda-forge |
| r-htmltools | 0.3.6 | r36he1b5a44_1003 | conda-forge |
| r-htmlwidgets | 1.3 | r36h6115d3f_1001 | conda-forge |
| r-httpuv | 1.5.1 | r36h0357c0b_1 | conda-forge |
| r-jsonlite | 1.6 | r36hcdcec82_1001 | conda-forge |
| r-labeling | 0.3 | r36h6115d3f_1002 | conda-forge |
| r-lambda.r | 1.2.3 | r36h6115d3f_1001 | conda-forge |
| r-later | 0.8.0 | r36h0357c0b_1 | conda-forge |
| r-lattice | 0.20_38 | r36hcdcec82_1002 | conda-forge |
| r-lazyeval | 0.2.2 | r36hcdcec82_1 | conda-forge |
| r-locfit | 1.5_9.1 | r36h516909a_1004 | conda-forge |
| r-magrittr | 1.5 | r36h6115d3f_1002 | conda-forge |
| r-mass | 7.3_51.4 | r36hcdcec82_1 | conda-forge |
| r-matrix | 1.2_17 | r36hcdcec82_1 | conda-forge |
| r-matrixstats | 0.54.0 | r36hcdcec82_1001 | conda-forge |
| r-memoise | 1.1.0 | r36h6115d3f_1002 | conda-forge |
| r-mgcv | 1.8_28 | r36hcdcec82_1 | conda-forge |
| r-mhsmm | 0.4.16 | r36h516909a_1003 | conda-forge |
| r-mime | 0.7 | r36hcdcec82_1 | conda-forge |
| r-munsell | 0.5.0 | r36h6115d3f_1002 | conda-forge |
| r-mvtnorm | 1.0_11 | r36h9bbef5b_1 | conda-forge |

| | | | |
|---|---|---|---|
| r-nlme | 3.1_141 | r36h9bbef5b_1 | conda-forge |
| r-optparse | 1.6.2 | r36h6115d3f_1 | conda-forge |
| r-permute | 0.9_5 | r36_1 | conda-forge |
| r-pillar | 1.4.2 | r36h6115d3f_2 | conda-forge |
| r-pkgconfig | 2.0.2 | r36h6115d3f_1002 | conda-forge |
| r-plogr | 0.2.0 | r36h6115d3f_1002 | conda-forge |
| r-plyr | 1.8.4 | r36h0357c0b_1003 | conda-forge |
| r-prettyunits | 1.0.2 | r36h6115d3f_1002 | conda-forge |
| r-promises | 1.0.1 | r36h0357c0b_1001 | conda-forge |
| r-r.methodss3 | 1.7.1 | r36h6115d3f_1002 | conda-forge |
| r-r.oo | 1.22.0 | r36h6115d3f_1001 | conda-forge |
| r-r.utils | 2.9.0 | r36h6115d3f_1 | conda-forge |
| r-r6 | 2.4.0 | r36h6115d3f_2 | conda-forge |
| r-rcolorbrewer | 1.1_2 | r36h6115d3f_1002 | conda-forge |
| r-rcpp | 1.0.2 | r36h0357c0b_0 | conda-forge |
| r-rcurl | 1.95_4.12 | r36hcdcec82_1 | conda-forge |
| r-reshape2 | 1.4.3 | r36h0357c0b_1004 | conda-forge |
| r-rlang | 0.4.0 | r36hcdcec82_1 | conda-forge |
| r-rsqlite | 2.1.2 | r36h0357c0b_0 | conda-forge |
| r-scales | 1.0.0 | r36h0357c0b_1002 | conda-forge |
| r-shiny | 1.3.2 | r36h6115d3f_1 | conda-forge |
| r-shinydashboard | 0.7.1 | r36h6115d3f_1001 | conda-forge |
| r-snow | 0.4_3 | r36h6115d3f_1001 | conda-forge |
| r-sourcetools | 0.1.7 | r36he1b5a44_1001 | conda-forge |
| r-stringi | 1.4.3 | r36h0357c0b_2 | conda-forge |
| r-stringr | 1.4.0 | r36h6115d3f_1 | conda-forge |
| r-tibble | 2.1.3 | r36hcdcec82_1 | conda-forge |
| r-upsetr | 1.4.0 | r36h6115d3f_1 | conda-forge |
| r-utf8 | 1.1.4 | r36hcdcec82_1001 | conda-forge |
| r-vctrs | 0.2.0 | r36hcdcec82_1 | conda-forge |
| r-viridislite | 0.3.0 | r36h6115d3f_1002 | conda-forge |
| r-withr | 2.1.2 | r36h6115d3f_1001 | conda-forge |
| r-xml | 3.98_1.20 | r36hcdcec82_1 | conda-forge |
| r-xtable | 1.8_4 | r36h6115d3f_2 | conda-forge |
| r-yaml | 2.2.0 | r36hcdcec82_1002 | conda-forge |
| r-zeallot | 0.1.0 | r36h6115d3f_1001 | conda-forge |
| ratelimiter | 1.2.0 | py36_1000 | conda-forge |
| readline | 8.0 | hf8c457e_0 | conda-forge |
| requests | 2.22.0 | py36_1 | conda-forge |
| rsa | 3.4.2 | py_1 | conda-forge |
| ruamel-yaml | 0.16.5 | pypi_0 | pypi |
| ruamel-yaml-clib | 0.1.2 | pypi_0 | pypi |
| s3transfer | 0.2.1 | py36_0 | conda-forge |
| samtools | 1.9 | h8571acd_11 | bioconda |
| sed | 4.7 | h1bed415_1000 | conda-forge |
| setuptools | 41.0.1 | py36_0 | conda-forge |
| simplejson | 3.16.1 | py36h470a237_0 | conda-forge |
| sip | 4.19.8 | py36hf484d3e_1000 | conda-forge |
| six | 1.12.0 | py36_1000 | conda-forge |
| smmap2 | 2.0.5 | py_0 | conda-forge |
| snakemake | 5.5.4 | 1 | bioconda |
| snakemake-minimal | 5.5.4 | py_1 | bioconda |
| spectra | 0.0.11 | py_1 | conda-forge |
| sqlite | 3.29.0 | hcee41ef_0 | conda-forge |
| tk | 8.6.9 | hed695b0_1002 | conda-forge |
| tktable | 2.10 | h555a92e_1 | conda-forge |
| toolshed | 0.4.6 | py_1 | bioconda |
| tornado | 6.0.3 | py36h516909a_0 | conda-forge |
| typing_extensions | 3.7.4 | py36_0 | conda-forge |
| urllib3 | 1.25.3 | py36_0 | conda-forge |
| wg-blimp | 0.9.3 | dev_0 | <develop> |
| wheel | 0.33.6 | py36_0 | conda-forge |
| wrapt | 1.11.2 | py36h516909a_0 | conda-forge |
| xmlrunner | 1.7.7 | py_0 | conda-forge |
| xorg-fixesproto | 5.0 | h14c3975_1002 | conda-forge |
| xorg-inputproto | 2.3.2 | h14c3975_1002 | conda-forge |
| xorg-kbproto | 1.0.7 | h14c3975_1002 | conda-forge |
| xorg-libice | 1.0.10 | h516909a_0 | conda-forge |
| xorg-libsm | 1.2.3 | h84519dc_1000 | conda-forge |
| xorg-libx11 | 1.6.8 | h516909a_0 | conda-forge |
| xorg-libxau | 1.0.9 | h14c3975_0 | conda-forge |
| xorg-libxdmcp | 1.1.3 | h516909a_0 | conda-forge |
| xorg-libxext | 1.3.4 | h516909a_0 | conda-forge |
| xorg-libxfixes | 5.0.3 | h516909a_1004 | conda-forge |
| xorg-libxi | 1.7.10 | h516909a_0 | conda-forge |
| xorg-libxpm | 3.5.12 | h14c3975_1002 | conda-forge |
| xorg-libxrender | 0.9.10 | h516909a_1002 | conda-forge |
| xorg-libxt | 1.1.5 | h516909a_1003 | conda-forge |
| xorg-libxtst | 1.2.3 | h14c3975_1002 | conda-forge |
| xorg-recordproto | 1.14.2 | h14c3975_1002 | conda-forge |
| xorg-renderproto | 0.11.1 | h14c3975_1002 | conda-forge |
| xorg-xextproto | 7.3.0 | h14c3975_1002 | conda-forge |
| xorg-xproto | 7.0.31 | h14c3975_1007 | conda-forge |
| xz | 5.2.4 | h14c3975_1001 | conda-forge |
| yaml | 0.1.7 | h14c3975_1001 | conda-forge |

| | | | |
|---|---|---|---|
| yarl | 1.3.0 | py36h14c3975_1000 | conda-forge |
| zlib | 1.2.11 | h516909a_1005 | conda-forge |
| zstd | 1.4.0 | h3b9ef0a_0 | conda-forge |
